# Nucleoprotein reassortment enhanced transmissibility of H3 1990.4.a clade influenza A virus in swine

**DOI:** 10.1101/2023.10.30.564250

**Authors:** Megan N. Thomas, Giovana Ciacci Zanella, Brianna Cowan, C. Joaquin Caceres, Daniela S. Rajao, Daniel R. Perez, Phillip C. Gauger, Amy L. Vincent Baker, Tavis K. Anderson

**Affiliations:** Department of Veterinary Microbiology and Preventive Medicine, College of Veterinary Medicine, Iowa State University, Ames, Iowa, USA; Department of Population Health, College of Veterinary Medicine, University of Georgia, Athens, GA, USA; Virus and Prion Research Unit, National Animal Disease Center, USDA-ARS, Ames, IA, USA

**Keywords:** influenza a virus, nucleoprotein, reassortment, viral transmission, reverse genetics

## Abstract

The increased detection of H3 C-IVA (1990.4.a) clade influenza A viruses (IAV) in U.S. swine in 2019 was associated with a reassortment event to acquire an H1N1pdm09 lineage nucleoprotein (pdmNP) gene, replacing a TRIG lineage NP (trigNP). We hypothesized that acquiring the pdmNP conferred a selective advantage over prior circulating H3 viruses with a trigNP. To investigate the role of the NP reassortment in transmission, we identified two contemporary 1990.4.a representative strains (NC/19 and MN/18) with different evolutionary origins of the NP gene. A reverse genetics system was used to generate wild-type (wt) strains and to swap the pdm and TRIG lineage NP genes, generating four viruses: wtNC/19-pdmNP, NC/19-trigNP, wtMN/18-trigNP, MN/18-pdmNP. Pathogenicity and transmission of the four viruses were compared in pigs. All four viruses infected 10 primary pigs and transmitted to 5 indirect contact pigs per group. Pigs infected via contact with MN/18-pdmNP shed virus two days earlier than pigs infected with wtMN/18-trigNP. The inverse did not occur for wtNC/19-pdmNP and NC/19-trigNP. These data suggest that reassortment to acquire a pdmNP gene improved transmission efficiency in the 1990.4.a, but this is likely a multigenic trait. Replacing a trigNP gene alone may not diminish the transmission of a wild-type virus sampled from the swine population. This study demonstrates how reassortment and subsequent evolutionary change of internal genes can result in more transmissible viruses that impact the detection frequency of specific HA clades. Thus, rapidly identifying novel reassortants paired with dominant HA/NA may improve prediction of strains to include in vaccines.

**Importance:** Influenza A viruses (IAV) are composed of eight non-continuous gene segments that can reassort during coinfection of a host, creating new combinations. Some gene combinations may convey a selective advantage and be paired together preferentially. A reassortment event was detected in swine in the United States that involved the exchange of two lineages of nucleoprotein (NP) genes (trigNP to pdmNP) that became a predominant genotype detected in surveillance. Using a transmission study, we demonstrated that exchanging the trigNP for a pdmNP caused the virus to shed from the nose at higher levels and transmit to other pigs more rapidly. Replacing a pdmNP with a trigNP did not hinder transmission, suggesting that transmission efficiency depends on interactions between multiple genes. This demonstrates how reassortment alters IAV transmission and that reassortment events can provide an explanation for why genetically related viruses with different internal gene combinations experience rapid fluxes in detection frequency.

## Introduction

The six internal genes of influenza A virus (IAV) contribute to pathogenesis and transmission in swine in ways that are less clearly defined than the surface genes hemagglutinin and neuraminidase (HA and NA). The internal genes encode the viral polymerase (PB2, PB1, PA), the nucleoprotein (NP), the matrix protein (M), and the non-structural protein (NS). IAV has a segmented genome and the ability to reassort, leading to different combinations of the eight gene segments and a vast potential for genetic diversity. Similar to the previously described requirement of balance between the HA and NA [1-3], there appear to be specific gene segments and/or combinations of internal genes, known as internal gene constellations, that may be preferentially selected among viral populations [4]. These combinations of internal genes are permissive to different HA and NA combinations and demonstrate sustained transmission in swine populations [5].

In swine, the internal genes are classified based on their evolutionary lineage: the triple reassortant lineage (TRIG), the human 2009 pandemic (pdm) lineage, or the live-attenuated influenza virus (LAIV) vaccine lineage. The internal gene constellation is abbreviated using the first letter of each lineage (T, P, or L) to form a six-character sequence in the following order: PB2, PB1, PA, NP, M, NS. The TRIG lineage emerged in the late 1990s and consists of PB1 gene segments derived from human seasonal H3N2; PB2 and PA gene segments from avian IAV; and NP, M, and NS gene segments from classical swine H1N1 [6-8]. Prior to the 2009 pandemic, all endemic viruses in the U.S. swine population contained the TRIG internal gene constellation, TTTTTT [5]. Following the H1N1 pandemic in 2009, the virus was repeatedly introduced into the pig population by humans [9-11]. The 8 gene segments in their entirety generally do not persist in U.S. pig populations for extended periods of time, but onward transmission of the pdm internal genes through reassortment was common [12]. From 2009 to 2012, the pdmM gene increased in abundance from 38% to 70%, making the TTTT<u>P</u>T constellation the second most commonly detected during this time period [13]. The pdmPA and pdmNP genes were also detected during this period but did not increase in abundance like the M gene [14-16].

In previous work, we demonstrated a rapid change in the detection frequency of the 1990.4.a clade of H3 IAV that began in 2019 [17]. Following the same general genome constellation patterns described above, the 1990.4.a internal gene constellation pre-pandemic was TTTTTT. The 1990.4.a viruses then acquired a pdmM gene, and this TTT<u>T</u>PT constellation became the most common in the clade between 2011 and 2018 [16]. However, the internal gene constellation found in the 1990.4.a major clade that expanded in detection frequency beginning in 2019 acquired a pdmNP gene [17]. Prior to 2019, this TTT<u>P</u>PT constellation was sporadically observed, but was uncommon (22 of 368 isolates detected between 2009 and 2016) [16].

The NP gene codes for a structural RNA-binding protein that, along with the trimeric polymerase complex and viral RNA, forms the ribonucleoprotein (RNP) particle. The NP is known to regulate multiple processes, including nuclear import/export of vRNP and transcription/replication [18]. In swine, results from a transmission study in pigs demonstrated that H3 viruses with the TTT<u>P</u>PT constellation were more effective in transmission when compared to H3 strains with a TTT<u>T</u>PT constellation [16]. Based on the function of the NP and the increasing detection of the TTT<u>P</u>PT constellation, we hypothesized that the success of the expanding clade of 1990.4.a H3 viruses could be explained by differences in the genetic lineage of the NP acquired following reassortment. We tested this hypothesis using a reverse genetics system to swap the respective pdm or TRIG lineage NP between two sustained 1990.4.a viruses. We demonstrated that the exchange of the pdmNP in the context of a genome previously containing a trigNP has the potential to improve transmission efficiency, but replacing a pdmNP with a trigNP does not necessarily have the opposite effect. Our results suggest viral transmission is dependent on multiple genes and the context of the other 7 genes of the genome.

## Methods

### Genetic analysis

In 2019, an NP reassortment event associated with increased detections of an H3 1990.4.a clade of viruses was detected [17]. There was a shift from a whole genome constellation of TTT<u>T</u>PT to the constellation including a pdmNP (TTT<u>P</u>PT) in the majority of viruses from this H3 HA clade. We collected all available complete whole genome sequences (WGS) of 1990.4.a viruses from 2016 to 2021 (n=177) from the USDA passive surveillance database, octoFLUshow [19], to quantify the frequency of detection of the internal gene constellations in this expanding clade over time. To determine detection frequency of the pdmNP in viruses of other HA clades, we also collated all complete swine H1 and H3 WGS from the USDA surveillance system (n=1720). The Nextstrain [20] platform was used to analyze the genetic diversity of the pdmNP genes (n=68) that paired with the H3 1990.4.a clade from 2010 to 2021. Augur commands translate and ancestral [21] were used to annotate amino acid substitutions on the backbone of the time-scaled maximum likelihood tree.

### Strain selection

Two IAV in swine viruses, A/swine/North Carolina/A02245294/2019 (NC/19) (Genbank accession numbers <u>MN646208.1</u>, <u>MN646209.1</u>, and <u>MT233822.1</u> - <u>MT233827.1</u>) and A/swine/Minnesota/A02266068/2018 (MN/18) (Genbank accession numbers <u>MK122759.1</u>, <u>MK122760.1</u>, and <u>MW308842.1 -</u> <u>MW308847.1</u>), were identified that had representative HA genes of the minor (TTT<u>T</u>PT) and major (TTT<u>P</u>PT) clades and were associated with the 1990.4.a expansion in 2019 (Figure 1) [17]. The remaining seven gene segments of these two strains were confirmed to represent the TTT<u>T</u>PT and TTT<u>P</u>PT constellations circulating in swine. First, we subsetted the larger 1990.4.a dataset described above to only include viruses with the internal gene constellations TTT<u>T</u>PT (n = 21) and TTT<u>P</u>PT (n=58) since 2018. Geneious Prime v2021.2 was used to construct two consensus sequences for each of the seven gene segments representing the TTTTPT and TTTPPT constellations. We used mafft v7.490 to confirm that the genes in NC/19 and MN/18 had at least 99.0% amino acid (AA) identity to the consensus sequence of the constellation they represented [22].

**Figure 1.**
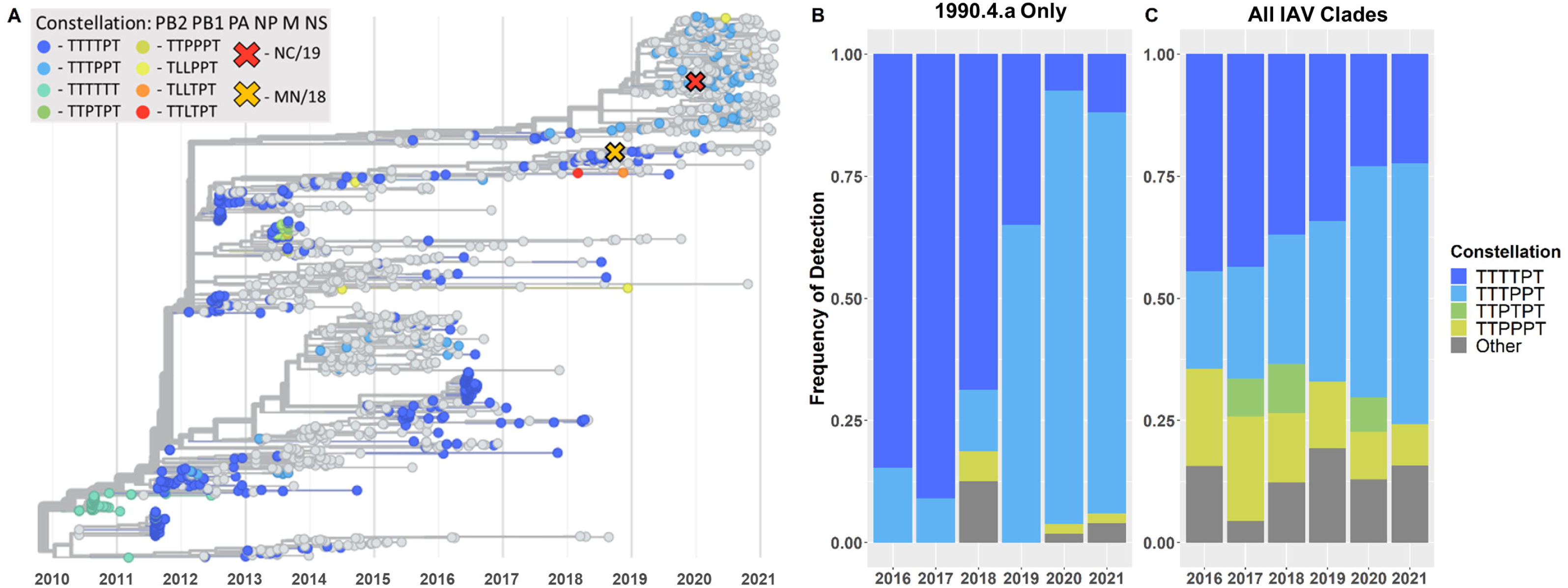
Summary of change in predominant internal gene constellation showed a shift from TRIG to pdm lineage NP over time. (A) Time-scaled phylogenetic tree of H3 1990.4.a IAV from 2010 to 2021. Chosen clade representative viruses, NC/19 and MN/18, are represented by an “X” on the tree. When possible, strains are colored by their internal gene constellation. (B) Frequency of detection of the most common internal gene constellations in H3 1990.4.a IAV from 2016 to 2021. (C) Frequency of detection of the most common internal gene constellation in all IAV clades from 2016 to 2021. The “Other” category in (B) and (C) includes any gene constellations not explicitly listed.

### Reverse genetics

Reverse genetics was used to generate four viruses using an 8-plasmid system as previously described [23, 24]. All the genes of the two representative viruses (NC/19 and MN/18) were synthesized (Twist Bioscience, San Francisco, CA) and subsequently cloned into the bidirectional plasmid vector pDP2002. For the transfection, co-cultures of 9×105 HEK293T and 1.5×105 MDCK cells were seeded per well in a 6-well plate. The following day, 1 µg of each plasmid was mixed with 18 µl of TransIT-LT1 transfection reagent (Mirus Bio LLC, Madison, WI). The mixture was incubated for 45 min and then used to overlay the 293T/MDCK cells overnight. The next day, the transfection mixture was replaced with fresh Opti-MEM media containing 1% AB (Life Technologies, Carlsbad, CA), and 24 h post-transfection, the media was supplemented with 1 µg/ml of tosylsulfonyl phenylalanyl chloromethyl ketone (TPCK) treated-trypsin (Worthington Biochemicals, Lakewood, NJ). Wild-type viruses were generated with their respective NP genes (wtNC/19-pdmNP and wtMN/18-trigNP). Reassortants of NC/19 and MN/18 were created by swapping only the NP genes between the two viruses (NC/19-trigNP and MN/18-pdmNP) and maintaining the remainder 7 genes from the respective parental strains. Viral sequences were confirmed by Sanger sequencing, and propagated in MDCK cells.

### *In vivo* animal study

A total of 65 healthy three-week-old, crossbred pigs were obtained from a herd free of IAV, PRRSV, and *Mycoplasma hyopneumoniae*. Upon arrival, pigs were confirmed to be seronegative to IAV NP antibodies by a commercial enzyme-linked immunosorbent assay (ELISA) kit (Swine Influenza Virus Antibody Test, IDEXX, Westbrook, ME). They were also treated with ceftiofur crystalline free acid and tulathromycin (Zoetis Animal Health, Florham Park, NJ) to reduce the risk of bacterial contaminants. Pigs were randomly selected for the non-challenge control (n=5) or four primary challenge groups (n=10/group), implanted with a Life Chip with Bio-Thermo Technology (Destron Fearing, DFW Airport, Texas), and moved to separate isolation rooms by group. The remaining pigs were assigned to be indirect contacts to the primary challenge pigs (n=5/group). All pigs were housed in biosafety level 2 (BSL2) containment and cared for in compliance with the Institutional Animal Care and Use Committee (IACUC) of the USDA National Animal Disease Center.

At approximately five weeks of age, the primary challenge pigs were inoculated with one of the four viruses: wtNC/19-pdmNP, NC/19-trigNP, wtMN/18-trigNP, MN/18-pdmNP. Pigs in the assigned group were inoculated intranasally (0.5 ml/nostril) and intratracheally (2 ml) with 1 x 10^5^ 50% tissue culture infectious dose (TCID_50_) per ml. For inoculation, pigs were anesthetized with an intramuscular injection of ketamine (8 mg/kg body weight; Phoenix, St. Joseph, MO), xylazine (4 mg/kg; Lloyd Inc., Shenandoah, IA), and tiletamine HCL/zolazepam HCL (Telazol; 6 mg/kg; Zoetis Animal Health, Florham Park, NJ). At 2 days post inoculation (DPI), indirect contact pigs were placed in separate raised decks in each of the four isolated challenge rooms. Nasal swabs (NS) and temperature readings were collected from the primary pigs on 0-5 DPI. NS were collected from contact pigs on 0-5, 7, and 9 days post contact (DPC). Nasal swabs were placed in tubes containing 2 ml minimum-essential-media (MEM) with 1 ul/ml L-1-tosylamido-2-phenylethyl chloromethyl ketone (TPCK) trypsin and stored at -80°C.

On 5 DPI, primary pigs were humanely euthanized with a lethal dose of pentobarbital sodium (Fatal Plus; Vortech Pharmaceuticals, Dearborn, MI) and necropsied, at which time the lungs were removed, scored for macroscopic lesions, and lavaged with 50ml of MEM containing 1% bovine serum albumin (BSA) to collect bronchoalveolar lavage fluid (BALF). BALF samples were plated on Casein and Blood Agar plates to evaluate for bacterial contamination prior to storing at -80°C. The distal trachea and right middle or affected lung lobe were collected and fixed in 10% buffered formalin, then processed using routine histologic methods to score microscopic lesions. On 11 DPC, contact pigs were bled and humanely euthanized using the same method described above.

### Virologic assays

Nasal swabs were filtered through a 0.2-micron syringe filter prior to virus isolation (VI). MDCK cells were grown to confluency in 48-well plates, which were then washed twice with phosphate-buffered saline (PBS). The filtered NS samples were then plated onto confluent MDCK cells and cultured at 37°C for 48 hours. Plates were fixed at 48 hours and stained via immunocytochemistry (ICC) as previously described [25]. To calculate a virus titer for NS that were positive by VI and all BALF samples, tenfold serial dilutions in MEM with 1:1000 TPCK trypsin were performed in triplicate onto confluent MDCK cells in 96-well plates. After 48 hours of incubation at 37°C, the plates were fixed and stained with the same protocol utilized for VI. Titers were calculated using the Reed and Muench method and TCID_50_/ml titers were log_10_ transformed for statistical analysis [26].

### Pathology

When the lungs were removed at necropsy, the percentage of lung surface affected by the purple-red consolidation typical of IAV infection was immediately recorded for each lobe. The weighted proportion of each lobe to the total lung volume was estimated to calculate the affected percentage of the entire lung [27]. The fixed lung and trachea sections were routinely processed and stained with hematoxylin and eosin. A veterinary pathologist blinded to the treatment groups scored the microscopic lesions using previously described parameters [28]. Parameters of each tissue were summed to calculate a total microscopic lesion score for the lung (0-22) and trachea (0-8).

### Serologic assays

Serum samples collected from the indirect contact pigs at 11 DPC were prepared for the hemagglutination inhibition (HI) assay by receptor-destroying enzyme (RDE II; Hardy Diagnostics, Santa Maria, CA) treatment, heat inactivation, and the addition of 50% turkey red blood cells to remove nonspecific hemagglutination inhibitors. HI assays were performed using test antigen from the respective wild-type challenge viruses with the same HA gene using previously described techniques [29].

### Statistical analysis

All analyses were performed using R Statistical Software (v4.0.5; R Core Team 2021). One-way analysis of variance followed by a Tukey-Kramer test were performed on the following variables using the core statistics package in R (v4.0.5; R Core Team): macroscopic pneumonia, microscopic lung and trachea lesions, and log_10_-transformed BALF and NS titers. Compact letter displays were generated when significant differences in group means (p-value <0.05) were present using the multcomp R package (v1.4.16; Bretz 2010).

## Results

### TTT<u>P</u>PT replaced TTT<u>T</u>PT as the majority IAV internal gene constellations in 2021

Since 2010, eight different internal gene constellations were paired with the H3 1990.4.a clade (Figure 1A), and the TTT<u>T</u>PT constellation comprised the majority of detections until 2019. There were sporadic TTT<u>P</u>PT detections during this timeframe, but only the detections after 2018 were sustained in the swine population. The TTT<u>P</u>PT constellation increased in frequency of detection in the H3 1990.4.a clade from 12.5% in 2018 to 65% in 2019 (Figure 1B). In 2020, the TTT<u>P</u>PT constellation further increased in detection making up 88.7% of the detections within the H3 1990.4.a clade. When evaluating all IAV H1 and H3 clades, the TTT<u>P</u>PT constellation detection frequency increased steadily, with 20.0% of detections in 2016 increasing to 53.3% of detections by 2021 (Figure 1C). The increased detection of the TTT<u>P</u>PT constellation was associated with a decreased detection of the TTT<u>T</u>PT constellation.

Two distinct clades of pdmNP were detected in the 1990.4.a clade between 2010 and 2021. The detections associated with the 2019 HA clade expansion were of the same lineage [17]. Three amino acid substitutions differentiate the lineage of pdmNP associated with the clade expansion from the lineage of previously detected pdmNP (I100V, K452R, and N377S). All three substitutions were estimated to have occurred between 2012 and 2014. There were no recent amino acid substitutions in the pdmNP associated with the increased detection of the TTT<u>P</u>PT constellation in the 1990.4.a clade beginning in 2019.

### Test strains represented contemporary detections and were significantly different

Two strains, NC/19 and MN/18, were chosen to represent the two contemporary H3 1990.4.a clades (Figure 1A). Both consisted of internal gene segments that were 99.0 to 100% identical at the amino acid level to the constellation consensus of the clade they were chosen to represent (Table 1). The two representative strains differed across their genomes, with gene segments ranging from 91.7 to 98.5% identical at the amino acid level (Table 1). The NP genes were 97.2% AA identical between NC/19 and MN/18, with their amino acid sequences differing at 14 positions (38, 70, 109, 217, 231, 301, 313, 353, 373, 377, 426, 456, and 473).

**Table 1.**
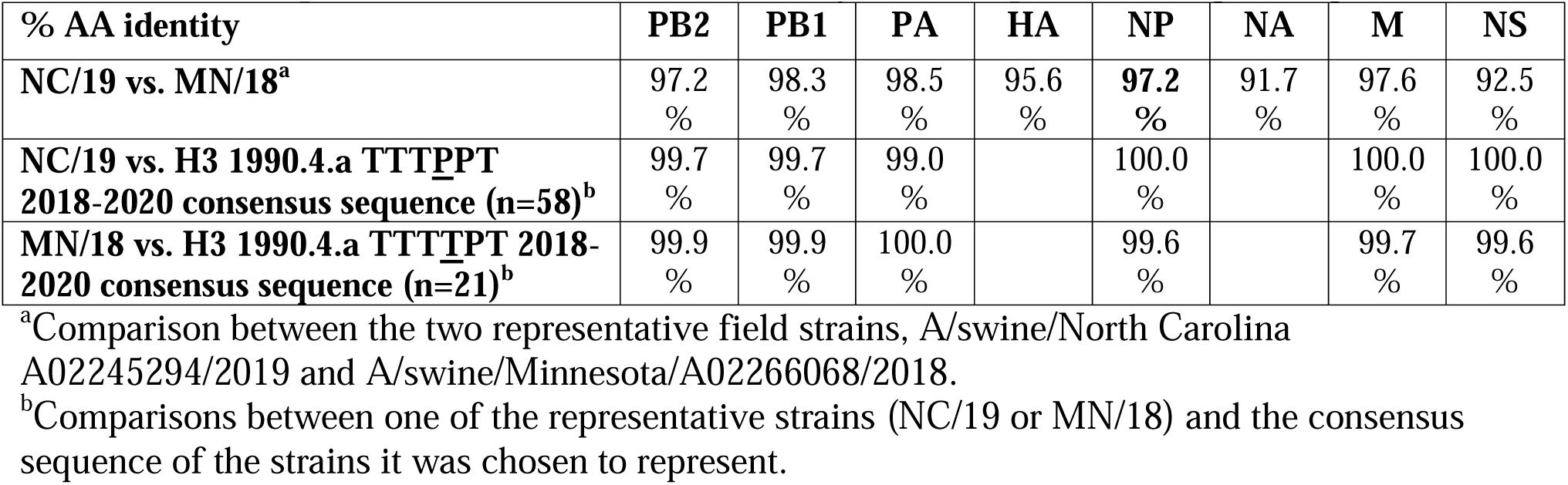
Percent pairwise amino acid (AA) identity for each pair of IAV gene segments.

### NP lineage did not affect pathology

All pigs infected with the four experimental strains selected to represent the NP reassortment exhibited limited to mild clinical signs consistent with experimental IAV infection in swine. The nonchallenged control pigs showed no clinical signs nor evidence of lung pathology indicative of IAV infection. All four of the challenge strains induced mild pneumonia in the primary pigs, with percentages of total lung affected that were not dependent on NP lineage (Table 2). There were no significant differences in mean microscopic lung or trachea scores between challenge groups (One-way ANOVA, p = 0.96, 0.80).

**Table 2.** Characteristics of infection with four challenge strains^a^.

| **Group** | **macroscopic pneumonia (%)** | **microscopic lung scores (0-22)** | **microscopic trachea scores (0-8)** | **BALF virus titer (log10 TCID50)** |
| --- | --- | --- | --- | --- |
| **Control** | 0.2 ± 0.1<sup>a</sup> | 0.0 ± 0.0<sup>a</sup> | 0.0 ± 0.0<sup>a</sup> | 0.0 ± 0.0<sup>a</sup> |
| **wtNC/19-pdmNP** | 6.4 ± 1.4<sup>c</sup> | 6.4 ± 0.7<sup>b</sup> | 3.3 ± 0.5<sup>b</sup> | 4.1 ± 0.3<sup>b</sup> |
| **NC/19-trigNP** | 5.0 ± 0.6<sup>bc</sup> | 6.3 ± 1.0<sup>b</sup> | 3.7 ± 0.4<sup>b</sup> | 5.4 ± 0.2<sup>c</sup> |
| **wtMN/18-trigNP** | 3.6 ± 0.6<sup>abc</sup> | 6.0 ± 0.6<sup>b</sup> | 3.2 ± 0.3<sup>b</sup> | 3.8 ± 0.4<sup>b</sup> |
| **MN/18-pdmNP** | 3.2 ± 0.5<sup>ab</sup> | 5.8 ± 1.1<sup>b</sup> | 3.2 ± 0.4<sup>b</sup> | 4.4 ± 0.2<sup>bc</sup> |

<sup>a</sup> Shown are the mean (± SEM) percentages of macroscopic lung consolidation, microscopic lung and trachea composite scores, and bronchoalveolar lavage fluid (BALF) virus titers of pigs experimentally inoculated with the challenge viruses or negative control pigs. Lowercase superscript letters depict statistically significant differences between values in the same column (p < 0.05).

### Acquisition of pdmNP improved transmission efficiency

IAV was isolated from the NS of all primary challenge pigs and the BALF of all but one pig, which was in the wtMN/18-trigNP challenge group. No virus was detected in the NS or BALF of nonchallenged negative control pigs. Nasal shedding of all primary challenged groups began on 1 DPI, decreased slightly on 2 DPI, and then was sustained at levels similar to 1 DPI until euthanasia on 5 DPI. The wtMN/18-trigNP challenge group had a trend for lower NS viral titers between 1-4 DPI, with a statistically significant difference compared to the other groups on 3 DPI (Figure 2; One-way ANOVA, p<0.001). The amount of virus in the BALF did not differ significantly between the two groups challenged with the two MN/18 viruses, but the NC/19-trigNP group had significantly higher BALF viral titers than the wtNC/19-pdmNP group (Table 3; One-way ANOVA, p = 0.50, p < 0.01).

**Figure 2.**
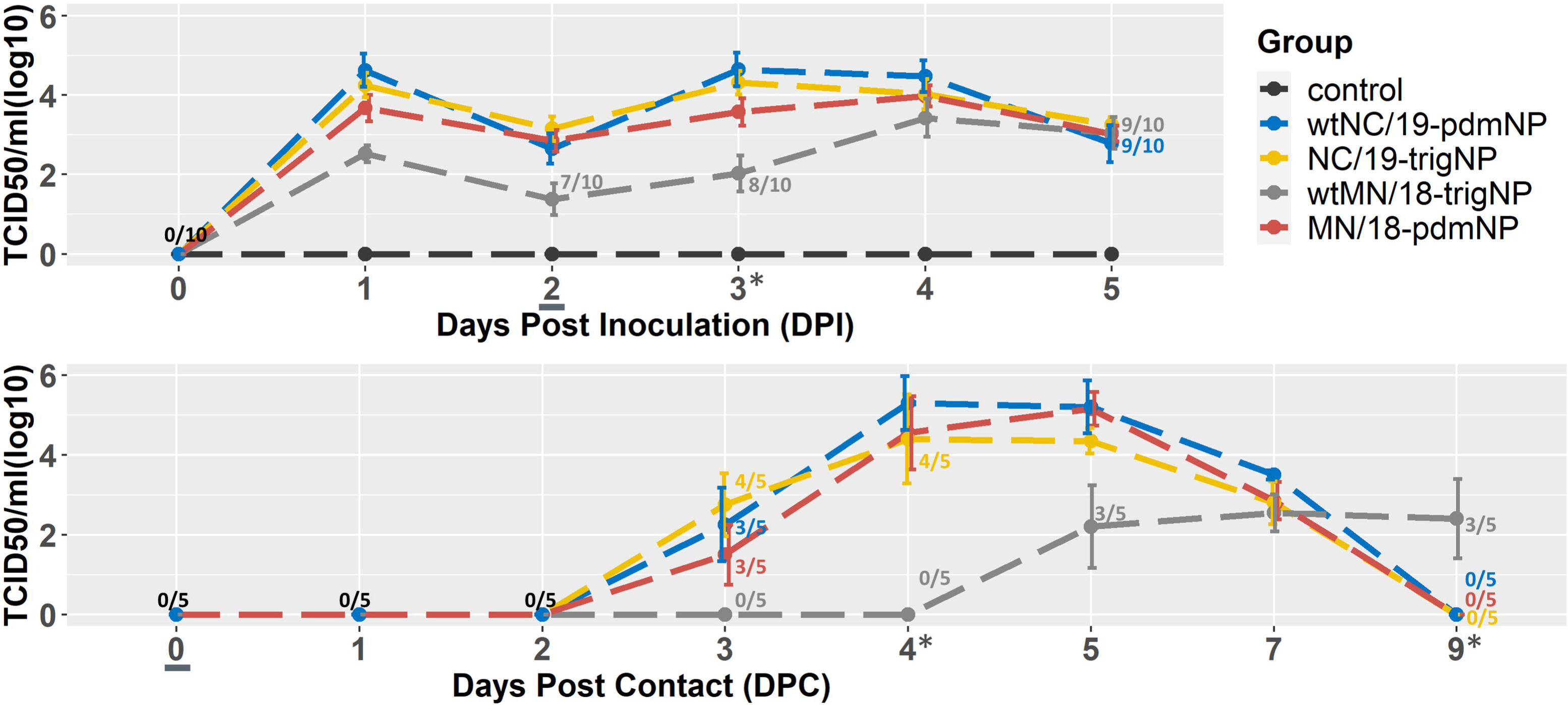
Mean (±SEM) viral titers over the (A) five days post inoculation for 10 primary pigs and (B) nine days post contact for 5 indirect contact pigs. The contacts were added on 2 DPI which corresponds with 0 DPC, denoted by an <u>underline</u>. Days when the viral titer of the wtMN/18-trigNP challenge group significantly differed from the other challenge groups are denoted by an asterisk (*). Days when <100% of pigs were shedding are labeled as n/10 (A) or n/5 (B) and colored by the group to which they belong. Black coloring indicates this number applies to all groups.

**Table 3.** Hemagglutination inhibition titers in serum from contact pigs at 11 DPC.

| **HI Assay** | **wtNC/19-pdmNP** |  |  |  |  | **NC/19-trigNP** |  |  |  |  |
| --- | --- | --- | --- | --- | --- | --- | --- | --- | --- | --- |
| **Pig** | 1 | 2 | 3 | 4 | 5 | 1 | 2 | 3 | 4 | 5 |
| **wtNC/19-pdmNP** | 160 | 160 | 80 | 160 | 160 | 640 | 160 | 160 | 80 | 160 |
|  | **wtMN/18-trigNP** |  |  |  |  | **MN/18-pdmNP** |  |  |  |  |
| **Pig** | 1 | 2 | 3 | 4 | 5 | 1 | 2 | 3 | 4 | 5 |
| **wtMN/18-trigNP** | 10 | 80 | 40 | 10 | 80 | 80 | 80 | 80 | 80 | 160 |

The indirect contact groups were added on 2 DPI/0 DPC; all four challenge viruses successfully transmitted to indirect contact pigs, i.e., virus was detected by titration from NS and BALF. Virus was detected in the NS of at least 3 out of 5 pigs in each group by 3 DPC (Figure 2); however, IAV was not detected in the wtMN/18-trigNP contact group NS until 5 DPC (3 out of 5). By 9 DPC, virus was no longer detectable in the NS of the wtNC/19-pdmNP, NC/19-trigNP, and MN/18-pdmNP contact pigs. At the same timepoint, 3/5 of the wtMN/18-trigNP contact pigs were still positive by NS but did not reach the same maximum mean group titer as the contacts in the other virus groups. These data suggest the transmission curve for the wtMN/19-trigNP was delayed by approximately two days along with a reduction in group mean titers in nasal secretions, while the MN/19-pdmNP transmission dynamics were comparable with the two NC/19 viruses.

### Reduced transmission efficiency delayed seroconversion

HI assays against the homologous virus were performed on the terminal serum of each indirect contact pig. Using a cutoff of HI titer ≥ 20, all indirect contact pigs seroconverted by the time of euthanasia at 11 DPC for the wtNC/19-pdmNP, NC/19-trigNP, and MN/18-pdm NP contact groups (Table 3). Two pigs (#622 and #625) in the wtMN/18-trigNP contact group did not seroconvert by this time point, and the HI titers of the wtMN/18-trigNP contact group were the lowest of the four groups.

## Discussion

The observed genetic diversity of IAV in swine has been substantially shaped by the emergence of the H1N1 pandemic in 2009 and subsequent human-to-swine transmission events. Following interspecies transmission, the internal genes of the human H1N1pdm09 virus reassort with endemic swine IAV, and these endemic swine IAV with H1N1pdm09-origin gene segments continue to diffuse through the swine IAV population. In 2011, the pdm lineage M gene became more commonly detected in IAV isolates from swine than the TRIG lineage M gene [30]. However, the majority of the other internal genes sequenced from swine isolates have remained of the TRIG lineage. We previously reported that H3 1990.4.a viruses first expanded the TTTPPT constellation in 2019 [17]. Here, we report that in 2021, twelve years after the 2009 pandemic, the majority of all H1 and H3 IAV in swine in the U.S. contained pdmNP genes. To investigate the expansion of the H3 1990.4.a clade, we used a reverse genetics approach to generate two wild-type viruses and then exchanged the pdm- and TRIG-lineage NP genes between them. We evaluated the pathogenesis and transmission of these four viruses in commercial weaned pigs. Our results showed that acquisition of the pdmNP improved transmission efficiency in the context of the MN/18 virus genome but replacing a pdmNP gene alone may not diminish transmission of an otherwise successful virus. Though this is a subtle difference, a virus that transmits 2 days sooner may “beat” other viruses to infection of the remainder of the susceptible population within a swine production system and effectively outcompete them, causing certain genes to achieve dominance in the population.

It was expected that all four viruses would infect the primary challenged pigs and successfully transmit to the indirect contacts as we demonstrated. We used two wild-type viruses that were representative of H3N2 viruses routinely isolated from field cases submitted for diagnostic investigation, indicating both viruses were fit for the swine host. Additionally, these two strains contained two of the most common internal gene constellations found in all H1 and H3 IAV circulating in swine in the USA, TTT<u>T</u>PT and TTT<u>P</u>PT. When compared to the wtNC/19-pdmNP virus, the wtMN/18-trigNP virus had decreased nasal shedding in the primary pigs at 1 DPI and 3 DPI, and infection of the indirect contact pigs was delayed by approximately 2 days. When the trigNP was replaced with a pdmNP in the MN/18 virus (MN/18-pdmNP), the magnitude of nasal shedding and transmission kinetics were comparable to the wtNC/19-pdmNP. Because the NP gene was the only difference between these two viruses, we conclude that acquisition of the pdmNP gene could improve the transmission efficiency for this virus. The NP protein is expressed in the early stage of infection and regulates multiple processes in the virus lifecycle, including nuclear import and export of viral RNP and transcription and replication of viral RNA [31, 32]. These roles in viral replication affect transmission efficiency by the impact on nascent virus production.

In contrast to the significant impact of the pdmNP on the MN/18 virus, nasal shedding and transmission to indirect contacts did not differ between the NC/19 viruses when the pdmNP was replaced with a trigNP. There were other differences across the genomes of the NC/19 and MN/18 viruses, such as those in the NA and NS genes, that could explain why the impacts of the NP were not bi-directional. Thus, it is likely that transmission efficiency is a multigenic trait. A parallel idea exists when we consider transmission of IAV within a new host species. Supporting this proposition was an *in vivo* pathogen and transmission study performed with reassortant viruses created to assess the contributions of gene combinations to replication and transmission of IAV in a novel host species [33]. This study used a human-seasonal H3 virus and a swine H3N1 virus and demonstrated that a swine-adapted HA gene alone could confer the ability to replicate in the lungs in the context of human NA and internal genes, but that the ability to replicate in the upper respiratory tract was dependent on the interaction between the HA gene and the NA or internal genes. Thus it is likely that transmission and viral phenotype relies upon individual gene segments and the interactions between all gene segments in the virus.

In our study, reassortment to acquire a pdmNP may not be the only mechanism explaining the 2019 H3N2 1990.4.a expansion. In our previous investigation, emergence of this clade was associated with the pdmNP reassortment, and a substitution in a key HA antigenic site (N156H) of the expanding H3 genes [17]. Additionally, population immunity to the diversity of H3 viruses in swine was also a likely contributor to the detection frequency of the H3 1990.4.a genetic clade [17]. Though we documented earlier transmission of the virus with the pdmNP, there was no statistically significant difference between the wtNC/19-pdmNP and NC/19-trigNP viruses in the parameters we measured. Further, the NC/19 representative of the expanding clade was capable of replicating and transmitting effectively in pigs and was robust to changes in the NP gene, i.e., inserting the trigNP did not diminish the transmission efficiency of the virus. These data further support the proposition that replication and transmission efficiency of IAV in the swine host, and likely virulence, are multigenic traits.

Genomic surveillance of influenza A virus in swine has been conducted systematically for over a decade in the US [19], but most of the effort has been guided towards analyzing the HA and, to a lesser extent, the NA gene(s). These surveillance data have been used to determine spatial and temporal trends in IAV in swine genetic diversity [15, 34, 35], support public health pandemic preparedness [8, 36], and has facilitated the selection of strains for use as antigens in vaccines [37-39]. However, our data reveals that understanding reassortment and the interactions between internal genes and the surface proteins is necessary to understand the dynamics of HA/NA clade detection frequency and turnover. Several veterinary diagnostic laboratories now offer whole genome sequencing for IAV, and the USDA IAV in swine surveillance system routinely generates whole genome sequences of swine IAV isolates. Based on current knowledge, the high genetic diversity and interplay between genes has made sequencing the entire viral genome a necessity. We show the impact that the NP gene can have on the transmission efficiency of an isogenic virus and hypothesize that the NP gene provided a competitive advantage to certain viruses. Routine whole genome sequencing can facilitate predicting when an HA/NA combination will increase or decrease in detection frequency through the identification of internal genes or constellations that have evidence for impact on transmission, or represent unique pairings, thus improving the efficacy and utility of monitoring and surveillance programs.

An unresolved question within our study is why it took twelve years for this pdmNP gene that has demonstrated a benefit to transmission to become dominant in the genomes of current IAV viruses in swine. When H1N1pdm2009 viruses were originally transmitted from humans into swine as a sequela of the pandemic, they rarely maintained purely H1N1pdm09 genomes: the surface proteins tended to persist when they underwent reassortment events that replaced human pdm internal genes with those endemic in swine [10, 40]. However, within two years the pdm M gene provided a selective advantage in the swine host and became the primary M gene in endemic IAV. The pdmNP described in this study has been maintained and evolving in the swine population following a human-to-swine spillover event in 2009-2010 influenza season [17]. Despite this early spillover, the pdmNP did not demonstrate significant changes in detection frequency like the pdm M gene and persisted at relatively low levels. Our data suggest the rapid increase also reflects the multigenic nature of IAV transmission: specifically, the pdmNP was only detected within the H3 1990.4.a clade at substantial levels beginning in 2019, and prior to the reassortment event, the pdmNP had acquired three amino acid substitutions. It is plausible that the impact of the pdmNP on transmission was only evident following the acquisition of these mutations, when paired with specific surface proteins, and under a specific immunological landscape reflecting vaccination and exposure to different cocirculating IAV in swine. IAV in swine are constantly evolving and reassortment to acquire divergent gene segments generates a vast amount of genetic diversity upon which selection can act to drive adaptation to the swine host and influence transmission and virulence. Future studies on the diversity and evolution of the pdmNP and implications of acquisition into swine IAV genomes are necessary to understand whether this transmission benefit is a general property of the gene, amino acids within the gene, or reflects specific gene pairings for IAV in swine.

## Data Availability

All sequence data are publicly available in NCBI GenBank, and the code, data, and supplementary material associated with this study are provided at https://github.com/flu-crew/datasets.

## Acknowledgments

We gratefully acknowledge pork producers, swine veterinarians, and laboratories for participating in the USDA Influenza A Virus in Swine Surveillance System and publicly sharing sequences. We thank USDA-ARS NADC animal resources unit caretakers and staff for assistance with animal studies and Celeste Snyder, Katharine Young, and Nicholas Otis for technical assistance. This work was supported in part by: the Iowa State University Presidential Interdisciplinary Research Initiative; the Iowa State University Veterinary Diagnostic Laboratory; the U.S. Department of Agriculture (USDA) Agricultural Research Service [ARS project number 5030-32000-231-000-D]; the National Institute of Allergy and Infectious Diseases, National Institutes of Health, Department of Health and Human Services [contract number 75N93021C00015 and 75N93021C00014]; the USDA Agricultural Research Service Research Participation Program of the Oak Ridge Institute for Science and Education (ORISE) through an interagency agreement between the U.S. Department of Energy (DOE) and USDA Agricultural Research Service [contract number DE-AC05-06OR23100]; and the SCINet project of the USDA Agricultural Research Service [ARS project number 0500-00093-001-00-D]. The funders had no role in study design, data collection and interpretation, or the decision to submit the work for publication. Mention of trade names or commercial products in this article is solely for the purpose of providing specific information and does not imply recommendation or endorsement by the USDA, DOE, ORISE, DARPA, or ISU. USDA is an equal opportunity provider and employer.

